# Clonal hematopoiesis is associated with increased toxicity in large B-cell lymphoma patients treated with chimeric antigen receptor T cell therapy

**DOI:** 10.1101/2021.09.28.461858

**Authors:** Neeraj Y. Saini, David M. Swoboda, Uri Greenbaum, Jungsheng Ma, Romil Patel, Kartik Devashish, Kaberi Das, Mark R. Tanner, Paolo Strati, Ranjit Nair, Luis E. Fayad, Sairah Ahmed, Hun Ju Lee, Swaminathan Iyer, Raphael Steiner, Nitin Jain, Loretta Nastoupil, Sanam Loghavi, Guilin Tang, Preetesh Jain, Michael Wang, Jason Westin, Michael R. Green, David Sallman, Eric Padron, Marco Davila, Frederick L. Locke, Richard Champlin, Elizabeth Shpall, Partow Kebriaei, Christopher R. Flowers, Michael Jain, Feng Wang, Andrew Futreal, Nancy Gillis, Sattva S. Neelapu, Koichi Takahashi

## Abstract

To explore the role of clonal hematopoiesis (CH) on chimeric antigen receptor (CAR) T-cell therapy outcomes, we performed targeted deep-sequencing on 114 large B-cell lymphoma patients treated with anti-CD19 CAR T-cells. We detected CH in 42 (36.8%) pre-treatment patient samples, most frequently in *PPM1D* (19/114) and *TP53* (13/114) genes. The incidence of grade ≥3 immune-effector cell-associated neurotoxicity syndrome (ICANS) was higher in CH-positive patients compared to CH-negative patients (45.2% vs. 25.0%, p=0.038). Higher toxicities with CH were primarily driven by three CH genes, *DNMT3A, TET2* and *ASXL1* (DTA mutations). The incidence of grade ≥3 ICANS [58.9% vs. 25%, p=0.02] and grade ≥3 cytokine release syndrome [17.7% vs. 4.2%, p=0.08] were higher in patients with DTA mutations than those without CH. The estimated 24-month cumulative incidence of therapy-related myeloid neoplasms after CAR-T therapy was higher in patients with CH than those without CH (19% [95%CI: 5.5-38.7] vs. 4.2% [95%CI: 0.3-18.4], p=0.028).

**Statement of Significance:** Our study reveals that clonal hematopoiesis mutations, especially those associated with inflammation (*DNMT3A, TET2, ASXL1*), are associated with severe grade toxicities in lymphoma patients receiving anti-CD19 chimeric antigen receptor therapy. Further studies to investigate the mechanisms and interventions to improve toxicities in the context of CH are warranted.

## Introduction

Adoptive T cell transfer therapy with chimeric antigen receptor (CAR)-T cells represents the latest breakthrough in the treatment of hematologic malignancies^1-3^. Three CD19-CAR-T cell products have received approval by regulatory medical agencies and were introduced into clinical practice for relapsed and refractory large B cell malignancies (r/r LBCL) (tisagenlecleucel, axicabtagene, and Iisocabtagene)^1-3^. Although durable responses have been observed in 30-40% of r/r LBCL patients treated with CAR-T therapy^1-3^, it is associated with significant systemic inflammatory toxicities such as cytokine release syndrome (CRS) and immune effector-cell associated neurotoxicity syndrome (ICANS) that are occasionally fatal^4^. Treatment-related toxicities with severe grade ≥3 CRS and/or ICANS occur in 10-31% of patients receiving CAR-T products^1-3^. Although there has been remarkable progress in the understanding and clinical management of CAR-T-related toxicities^5^, a significant knowledge gap exists in the mechanisms and host factors impacting these toxicities.

Clonal hematopoiesis (CH) is a clonally expanded population of hematopoietic stem cells bearing somatic gene mutations^6^. CH has been recognized as a driver of systemic inflammation^7^ and is associated with an increased risk of therapy-related myeloid neoplasms (t-MN) after chemotherapy^8,9^. Murine studies suggest that knockout of CH genes (*Dnmt3a* or *Tet2*) can contribute to a dysregulated inflammatory microenvironment by altering T–cell function^10,11^. Furthermore, recent clinical evidence indicates an emerging role of CH in accelerating graft versus host disease (GvHD) after allogeneic stem cell transplantation^12^. Since anti-CD19 CAR-T cell therapy, a highly effective therapy for LBCL and other lymphoid malignancies^13^, is associated with systemic inflammatory toxicities and given CH’s role in driving systemic inflammation, we hypothesized that CH influences the incidence and severity of CAR-T therapy toxicities. This study aimed to identify the clinical impact of CH in r/r LBCL patients undergoing CAR-T cell therapy.

## Results

### Patient characteristics and incidence of CH mutations

A total of 114 r/r LBCL patients at two different institutions, MD Anderson Cancer Center (MDACC, USA, n=99) and Moffitt Cancer Center (USA, n=15), whose peripheral blood (PB) buffy coat samples were available for CH analysis, were studied. The patient characteristics of the study cohort are listed in **Table 1**. Of the 114 patients with r/r LBCL, 105 were treated with axicabtagene cilolecleucel and 9 received tisagenlecleucel. The median age for the entire cohort was 63.0 years (range: 29.0 – 87.0 years) and patients received a median of 3 lines of systemic therapy prior to CAR-T therapy. The histological diagnosis was subclassified into DLBCL/high-grade B-cell lymphoma (n=91), transformed follicular lymphoma (n=21), and primary mediastinal lymphoma (n=2).

**Table 1.**
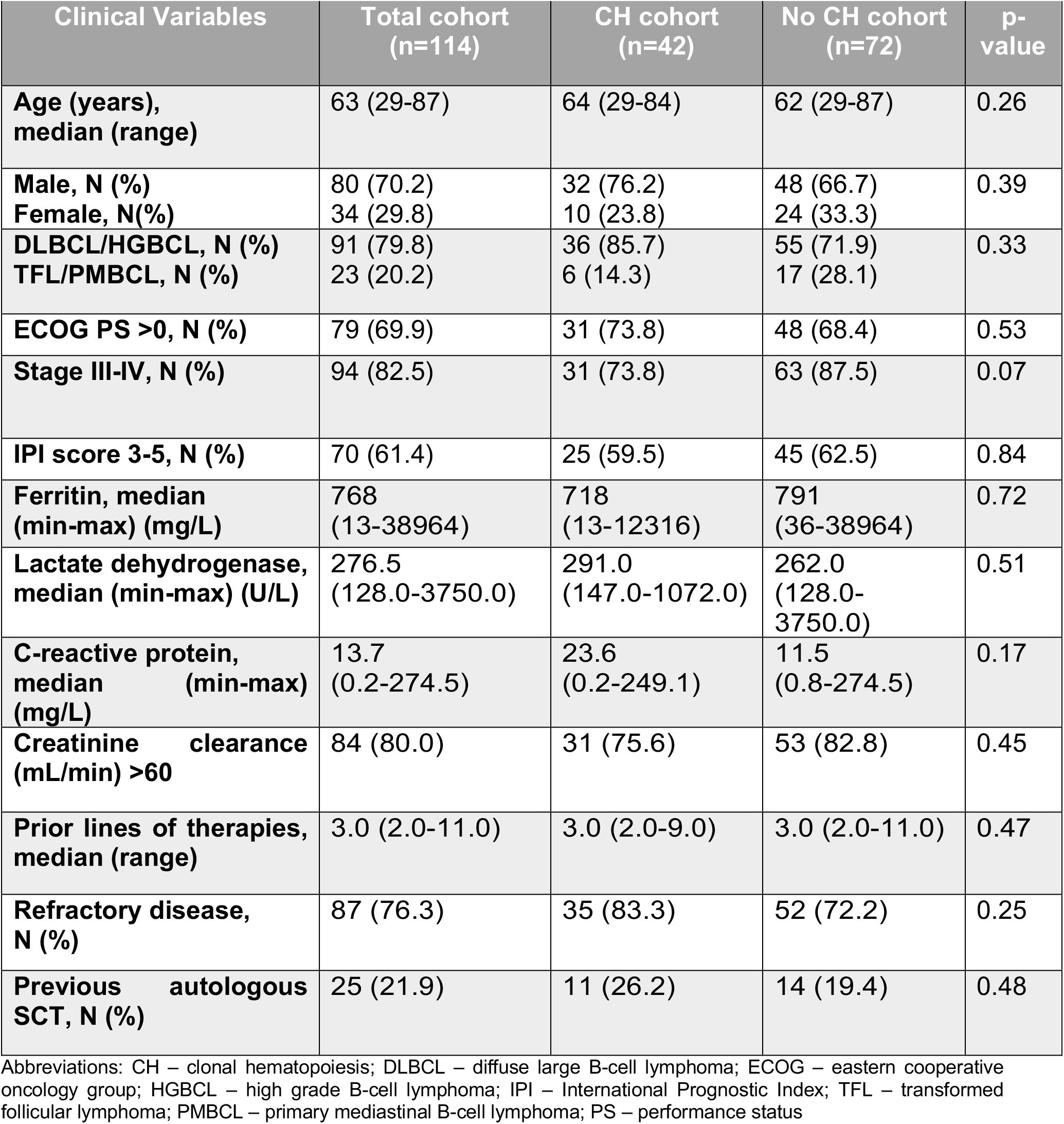
Baseline (day-5) characteristics of combined LBCL patients with anti-CD19 CAR-T therapy in LBCL patients.

| Clinical Variables | Total cohort<br>(n=114) | CH cohort<br>(n=42) | No CH cohort<br>(n=72) | p-value |
| --- | --- | --- | --- | --- |
| Age (years),<br>median (range) | 63 (29-87) | 64 (29-84) | 62 (29-87) | 0.26 |
| Male, N (%) | 80 (70.2) | 32 (76.2) | 48 (66.7) | 0.39 |
| Female, N(%) | 34 (29.8) | 10 (23.8) | 24 (33.3) |  |
| DLBCL/HGBCL, N (%) | 91 (79.8) | 36 (85.7) | 55 (71.9) | 0.33 |
| TFL/PMBCL, N (%) | 23 (20.2) | 6 (14.3) | 17 (28.1) |  |
| ECOG PS >0, N (%) | 79 (69.9) | 31 (73.8) | 48 (68.4) | 0.53 |
| Stage III-IV, N (%) | 94 (82.5) | 31 (73.8) | 63 (87.5) | 0.07 |
| IPI score 3-5, N (%) | 70 (61.4) | 25 (59.5) | 45 (62.5) | 0.84 |
| Ferritin, median<br>(min-max) (mg/L) | 768<br>(13-38964) | 718<br>(13-12316) | 791<br>(36-38964) | 0.72 |
| Lactate dehydrogenase,<br>median (min-max) (U/L) | 276.5<br>(128.0-3750.0) | 291.0<br>(147.0-1072.0) | 262.0<br>(128.0-3750.0) | 0.51 |
| C-reactive protein,<br>median (min-max)<br>(mg/L) | 13.7<br>(0.2-274.5) | 23.6<br>(0.2-249.1) | 11.5<br>(0.8-274.5) | 0.17 |
| Creatinine clearance<br>(mL/min) >60 | 84 (80.0) | 31 (75.6) | 53 (82.8) | 0.45 |
| Prior lines of therapies,<br>median (range) | 3.0 (2.0-11.0) | 3.0 (2.0-9.0) | 3.0 (2.0-11.0) | 0.47 |
| Refractory disease,<br>N (%) | 87 (76.3) | 35 (83.3) | 52 (72.2) | 0.25 |
| Previous autologous<br>SCT, N (%) | 25 (21.9) | 11 (26.2) | 14 (19.4) | 0.48 |

Abbreviations: CH – clonal hematopoiesis; DLBCL – diffuse large B-cell lymphoma; ECOG – eastern cooperative oncology group; HGBCL – high grade B-cell lymphoma; IPI – International Prognostic Index; TFL – transformed follicular lymphoma; PMBCL – primary mediastinal B-cell lymphoma; PS – performance status

CH was detected in the pre-treatment samples of 42 of the 114 (36.8%) patients. The complete list of genes and variants is provided in **Table S1**. The lab parameters on day - 5 prior to induction chemotherapy, including serum inflammatory markers, such as ferritin and C-reactive protein, were not significantly different between patients with and without CH (**Table 1, Figure S1**). The most frequently mutated genes were *PPM1D* (19/114, 16.7%), followed by *TP53* (13/114, 11.4%), *DNMT3A* (7/114, 6.1%), *TET2* (6/114, 5.2%) and *ASXL1* (4/114, 3.5%) (**Figures 1 and S2**). A total of 72 CH variants were detected in 42 patients with the median variant allele frequency (VAF) of CH of 5.8% (range: 2.1% - 49.5%) (**Figure S3**) and 19 variants in 15 patients were present at a VAF greater than 10%. In 30 (71.4%) patients, a single gene mutation was detected as CH, while 12 (28.6%) patients carried two or more gene mutations. The high proportion of patients having CH mutations in DNA damage pathway genes (*PPM1D* and *TP53)* was notable in this cohort (**Table S1**) and is likely associated with prior exposure to chemotherapies^14-17^. Among the 12 patients with more than one mutation, the most frequent combination was *TP53 and PPM1D* (n=8, 44.4%) mutations.

**Figure 1.**
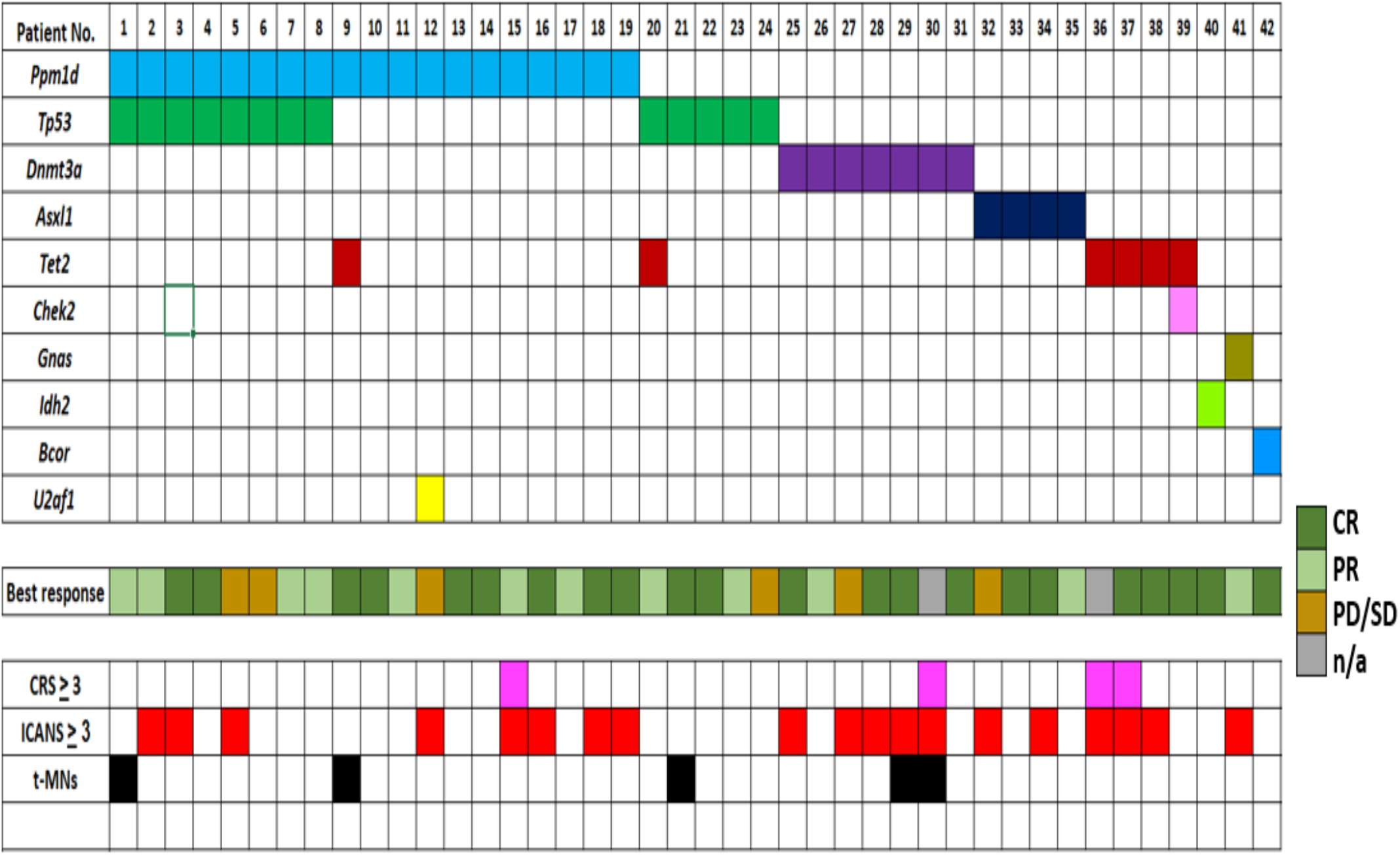
Oncoplot of clonal hematopoiesis (CH) mutations at baseline in LBCL patients treated with anti-CD19 CAR-T therapy and their association with response and toxicity outcomes. Abbreviations: CR – complete response; CRS – cytokine release syndrome; ICANS – immune cell associated neurotoxicity syndrome; n/a – not evaluable; t-MNs – treatment related myeloid neoplasms; PR – partial response; PD/SD – progressive disease/ stable disease.

### CH does not affect treatment response and survival outcomes with CAR-T therapy

The median duration of follow-up among survivors in our cohort was 14.9 (range: 1.2-30.5) months. The best overall response rate (ORR) and complete response (CR) for the whole cohort was 78.5% (84/107) and 56.1% (60/107), respectively. The rates of CR and ORR were not significantly different between patients with CH and without CH (CR: 55.0% vs. 56.7%, p=1.00, ORR: 85.0% vs. 74.6%, p=0.23, **Figure 2A**). The median progression-free survival (PFS) and overall survival (OS) for the whole cohort were 4.8 and 15.7 months, respectively (**Figure S4**). We did not observe any significant differences in PFS and OS between patients with CH and those without CH (**Figure 2B and S5**).

**Figure 2.**
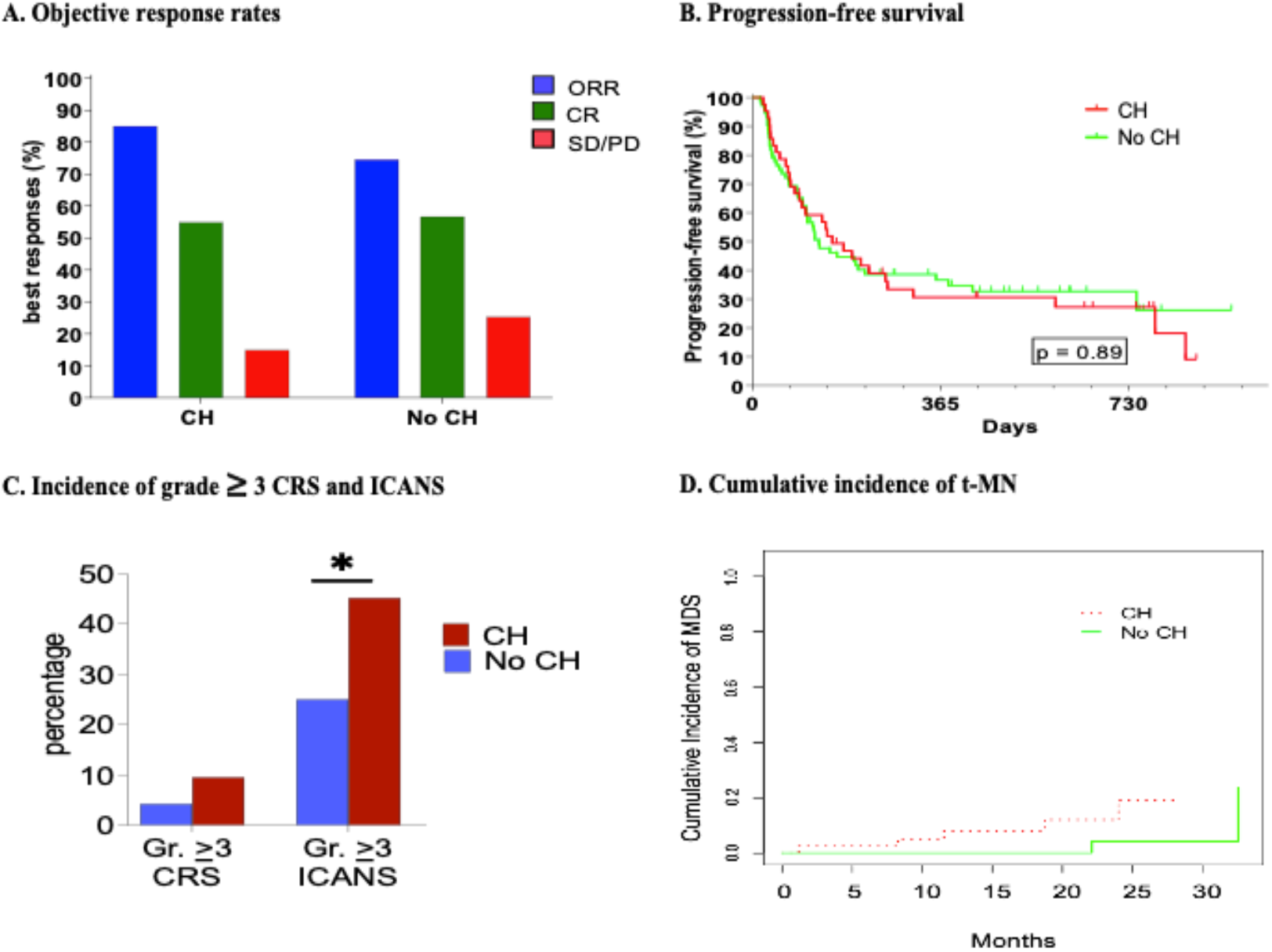
Associations between clinical outcomes and clonal hematopoiesis (CH) status in LBCL patients treated with anti-CD19 CAR-T therapy. A) Bar graph showing best response rates in CH versus no CH patients. B) Kaplan-Meier curve of progression-free survival in patients with CH and no CH. C) Bar graph showing incidence of grade 3/4 severe CRS and grade 3/4 severe ICANS in the CH and no CH patients. D) Cumulative incidence of therapy-related myeloid neoplasms in patients with CH compared to no CH. *p= 0.028. Abbreviations: CR – complete response; CRS – cytokine release syndrome; ICANS – immune cell associated neurotoxicity syndrome; ORR – overall response rates; t-MNs – treatment-related myeloid neoplasms; PR – partial response; PD – progressive disease; SD - stable disease.

### CH increases the risk of severe CAR-T related toxicities - CRS/ICANS

We also analyzed the impact of CH on CAR-T-associated toxicities. A total of 39 (92.9%) and 65 (90.3%) patients developed CRS with all grades in the CH and no-CH cohorts, respectively (p=0.743). A total of 24 (57.1%) and 37 (51.4%) developed ICANS with all grades in CH and no-CH cohorts, respectively (p=0.566). As we observed no differences in the incidence of all grades CRS or ICANs between the two cohorts, we next analyzed the incidence of severe toxicities (grades ≥3). There were 7 (6.1%) and 37 (32.5%) patients who had grade ≥3 CRS and grade ≥3 ICANS, respectively, in the entire population. While the overall incidence of grade ≥3 CRS was low in our cohort (6.1%), the incidence was numerically higher, but not statistically significant, in patients with CH (9.5%, 4/42) compared to the patients without CH (4.2%, 3/72) (p=0.42, **Figure 2C**). The rate of grade ≥3 ICANS was significantly higher in patients with CH, at 45.2% (19/42), compared to 25.0% (18/72 patients) in patients without CH (p=0.038, **Figure 2C**). On a multivariate analysis, the presence of CH was the only covariate significantly associated with an increased risk of grade ≥3 ICANS (odds ratio=2.47, 95% CI: 1.02-6.02, p=0.046, **Supplementary Table S4 and S5**). The percentage of patients requiring tocilizumab and corticosteroids for management of CRS and ICANS was comparatively higher, although not statistically significant, in patients with CH at 64.3% (27/42) and 52.4% (22/42), respectively, compared to 55.6% (40/72, p=0.43) and 43.1% (31/72, p=0.43) of patients with no-CH, respectively (**Figure S4**).

### Individual CH mutations have differential impact on CAR-T toxicity

We further analyzed the survival and toxicity outcomes associated with CH mutations that have been associated with inflammation in the literature, namely *DNMT3A, TET2*, and *ASXL1* (DTA mutations). In patients harboring DTA CH mutations, the incidence of grade ≥2 [70.5% (12/17) vs. 41.7% (30/72), p=0.06] or ≥3 [58.9% (10/17) vs. 25% (18/72), p=0.02] ICANS was significantly higher compared to in patients with no CH mutations (**Table 2**). Similarly, we saw a trend of increased grade ≥3 CRS in patients with DTA compared to patients with no CH mutations [17.7% (3/17) vs. 4.2% (3/72), p=0.08]. However, we did not find any difference in response rates between patients with DTA CH mutations or without CH mutations, as shown in **Table 2**.

**Table 2.** Survival and toxicity outcomes between patients with and without clonal hematopoiesis.

| Variables | No CH mutations | CH mutations | DTA CH mutations | p-value |  |
| --- | --- | --- | --- | --- | --- |
|  |  |  |  | CH vs. No CH | DTA CH vs. No CH |
| ICANS ≥ 2 | 30/72<br>(41.66%) | 22/42<br>(52.3%) | 12/17<br>(70.5%) | 0.33 | 0.056 |
| ICANS ≥ 3 | 18/72<br>(25%) | 19/42<br>(45.2%) | 10/17<br>(58.9%) | 0.04 | 0.02 |
| CRS ≥ 2 | 33/72<br>(45.8) | 20/42<br>(47.6%) | 9/17<br>(52.9%) | 1.0 | 0.79 |
| CRS ≥ 3 | 3/72<br>(4.2%) | 4/42<br>(9.5%) | 3/17<br>(17.7%) | 0.42 | 0.08 |
| CR | 38/67<br>(56.7%) | 22/40<br>(55%) | 10/15<br>(66.7%) | 1.0 | 0.57 |

### Plasma cytokine evaluations post-CAR-T infusion between CH and no-CH cohort

In patients with available samples, we also analyzed inflammatory cytokine levels in plasma at serial timepoints from day 0 until 2 weeks post-CAR-T infusion (n=43). There was a trend of higher median plasma levels of IL-6 at day 0 in CH patients (1.12 pg/ml) compared to no-CH patients (0.62 pg/ml) (p=0.058, **Table S6**), however, no differences were seen in other inflammatory cytokines. Also, we did not observe statistically significant differences in peak plasma levels of any inflammatory cytokines between CH and no-CH patients (**Table S7**).

### CH leads to increased therapy-related myeloid neoplasms

We assessed for cytopenia at leukapheresis and at day 90 post-CAR-T infusion in both the CH and no-CH cohorts. There were no differences in hemoglobin levels, platelet counts, and absolute lymphocyte and neutrophil counts at the time of leukapheresis or at day 90 post-CAR-T infusion (**Table S8**). We also compared the incidence of therapy-related myeloid neoplasms (t-MN) after CAR-T therapies. Seven patients developed t-MN following CAR-T therapy; 5 (5/42, 11.9%) had baseline CH and 2 (2/72, 2.8%) did not. At 24 months, the estimated cumulative incidence rates of t-MN after CAR-T therapy were 19% (95% CI: 5.5 - 38.7%) and 4.2% (95% CI: 0.3 - 18.4%) for patients with and without CH, respectively (p=0.028, **Figure 2D**). The clinical history and mutation analysis of the 5 patients with CH who subsequently developed t-MN are presented in **Table S9**. Mutational analyses were available in only 3 patients and among these, mutations were shared between CH and t-MN in two patients. The VAF levels of CH mutations on the diagnostic bone marrow samples corresponded to the blasts burden in these two patients. Information about these mutations are presented in **Table S9**.

## Discussion

In this cohort of heavily pretreated LBCL patients, we found that CH is associated with increased severe immune-mediated toxicities, particularly ICANS, following CAR-T cell therapy. Also, we found that CH did not impact survival outcomes or responses following CAR-T therapy. These findings add to the growing body of evidence linking CH with systemic inflammation in multiple clinical contexts such as atherosclerosis^18^, graft-versus-host-disease^12,19^, and infection^20^.

The incidence of CH in our cohort was approximately 40% and was similar to the recently reported incidence of CH (48%) in a mixed population of patients with lymphoma and myeloma undergoing CAR-T therapy at Dana-Farber Cancer institute (DFCI)^15^. These incidences are higher compared to that observed in LBCL patients undergoing autologous stem cell transplant (ASCT, CH incidence 25-30%)^14,17^. As a majority of the patients undergoing CAR-T were previously treated with ASCT, it is likely that there is a stepwise increase in the incidence of therapy-related CH with iterative exposures to chemotherapies. What was also notable in this cohort, as well as in the other heavily-treated cohorts, is the preponderance of CH with DNA damaging pathway genes, such as *PPM1D* and *TP53* mutations, which often co-occurred in the same patient. While it is difficult to dissect the clonal relationship of these two co-occurring mutations, our previous study using single-cell analysis indicated the mutually exclusive relationship of the two mutations at the cellular level ^21^.

In their study, the DFCI group reported an increase in CAR-T associated toxicities in the CH cohort, as well as higher CR rates^15^. There was a statistically significant increase in grade ≥2 CRS (77.8% CH vs. 45.9% no CH, p=0.042), albeit only in patients with age <60 years. Moreover, the incidence of grade ≥2 ICANS was comparatively much higher in CH patients (60% vs. 43%, p=0.06), although not statistically significant. These findings contrast with our results where we did not see a significant correlation between CH and CRS. One of the reasons could be the differences in the mutational pattern and frequencies in the two cohorts. Each CH mutation is biologically different and CH-harboring myeloid cells might spread in the tumor microenvironment differently^22^, leading to unique interactions with CAR-T cells and result in disparate toxicity outcomes. The DFCI cohort had a much higher number of *DNMT3A* and *TET2* mutations compared to our cohort and these mutations are well known in the literature to be associated with inflammation^23^. In fact, most of the high-grade toxicities in our cohort were driven by *DNMT3A/TET2/ASXL1* mutations; grade ≥3 CRS and grade ≥3 ICANS rates with these three mutations were 17.6% (3/17) and ∼60% (10/17), respectively. These subtle differences in the mutation spectrum might drive the discordant results seen in the DFCI study, both from toxicities as well as response perspectives. However, similar to our study, they did not observe any differences in PFS and OS between the two cohorts.

In contrast to CRS, where anti-IL-6 therapy has been shown to mitigate the development of severe CRS, there are currently no interventions available to prevent the development of severe ICANS. Consistent with this practice and the notion that CH is associated with systemic inflammation, we observed a statistically significant association between CH and severe ICANS. CH has a potential to influence outcomes and toxicity in CAR-T therapy through multiple mechanisms. In the tumor microenvironment, the severity of toxicities could be influenced by crosstalk between CH mutant myeloid cells, tumor cells, and CAR-T cells. This crosstalk could also influence the activation of bystander immune cells and lymphocytes in the tumor milieu, potentially leading to more inflammation and toxicities. Moreover, these CH mutants are associated with differential metabolism requirements and can produce micro changes in the metabolic signatures associated with tumor stroma^24^. In our cohort, we did not see a difference in inflammatory cytokines, either at baseline or at peak post CAR-T infusion, between the CH and no-CH cohorts. However, the cytokine repertoire we examined did not include IL-1, which has been shown to be well-associated with CH, especially the *TET2* mutation^23^. Moreover, IL-1 is strongly associated with ICANS pathophysiology^5^ and therefore, it will be important to explore the contribution of IL-1 in CH associated CAR-T toxicities.

In agreement with prior reports,^8,9,14^ we observed an increased rate of t-MNs in patients with CH receiving induction chemotherapy prior to CAR-T cell infusion, which portends poor outcomes. It is quite possible that through a longer follow-up and with a larger cohort, we might see poor outcomes in the CH cohort that is driven by a higher incidence of t-MNs, as seen in transplant settings^14^. Taken together, our results suggest that further studies are needed to elucidate the biological mechanisms by which CH influences immune-mediated toxicities associated with CAR-T cell therapy. Understanding the mechanisms by which CH influences toxicities may lead to novel intervention strategies to prevent high-grade CRS and ICANS after CAR-T therapy.

## Methods

### Patients and samples

Cryopreserved peripheral blood buffy coat samples from patients with r/r LBCL receiving standard of care axicabtagene ciloleucel (axi-cel) or tisagenlecleucel CAR-T therapy collected from the time of apheresis to any timepoints prior to induction chemotherapy were analyzed for CH detection. The MDACC cohort (n=99) consisted of consecutive LBCL patients who underwent anti-CD19 standard of care CAR-T therapy between 10/2018 to 6/2020 and whose frozen buffy coats were available in the Lymphoma Tissue Bank. Similarly, the Moffitt cohort included standard of care consecutive CAR-T patients whose peripheral blood was available for analysis. All patients provided written consent through an institutional review board-approved protocol at either The University of Texas MD Anderson Cancer Center (N = 99) or at the Moffitt Cancer Center (N = 15).

### DNA sequencing and bioinformatics pipelines to detect CH

The complete descriptions of sequencing and bioinformatics pipelines identifying high-confidence somatic single-nucleotide variants and indels from targeted capture DNA sequencing is provided in the supplement. In brief, for the MDACC cohort, the pre-treatment buffy coat samples were sequenced using a SureSelect custom panel of 300 genes (Agilent Technologies, Santa Clara, CA) that covers genes recurrently mutated in CH and hematologic malignancies. For the Moffitt cohort, DNA was extracted from peripheral blood for library preparation using a custom 76-gene hybrid-capture panel with unique molecular barcodes. We used a minimum variant allele frequency (VAF) cut-off of 2% for CH mutations, in accordance with a prior report^6^.

### Cytokine measurements

Available frozen plasma samples from the patients were obtained at different time-points from day 0 to day 14 and cytokines were measured using multiplex assays on a Meso-Scale Discovery platform^25^. The cytokines that were measured included IL-2, IL-4, IL-5, IL-16, IL-10, IL-13, IL-17A, granulocyte-macrophage colony-stimulating factor (GM-CSF), tumor necrosis factor-alpha (TNF-a) and interferon-gamma (IFN-g).

### Statistical analysis

Categorical covariates were summarized by frequencies and percentages and continuous covariates were summarized by means, standard deviations, medians, and ranges. Box-and-whisker plots were also used to summarize continuous variables. Comparisons between cohorts were performed using Fisher’s exact tests for categorical variables and Wilcoxon rank sum tests for continuous variables. A multivariable logistic regression model was fitted to evaluate associations between CH and ICANS adjusting covariates of interest. Unadjusted survival distributions were estimated by the Kaplan-Meier method and comparisons were made with the log rank test. Univariate Cox proportional hazards regression models were used to evaluate the associations between survival outcomes and the covariate of interest. The outcome variable of t-MNs was analyzed using competing risk models, where the competing risk was death. Gray’s test was used for comparisons of t-MNs between cohorts. PFS was defined as the time from the date of CAR-T infusion to the progression of disease or death or last follow-up (whichever occurred earlier). OS was defined as the time from the date of CAR-T cell infusion to death or last follow-up (whichever occurred earlier). A p-value of <0.05 (two-tailed) was considered statistically significant. Statistical analyses were conducted using R 3.6.1 and graph-pad PRISM 9 software.

## Acknowledgments

This work was supported in part by the University of Texas MD Anderson Cancer Center B-cell Lymphoma Moonshot (SSN), AML and MDS Moonshot (KT), Sabin Family Fellow Award (KT), American Society of Hematology Scholar Award (KT), Lyda Hill Foundation (AF), Physician Scientist Program at MD Anderson (KT), NIH/NCI R01 CA237291 (KT), and NCI Cancer Center Support Grant to the University of Texas MD Anderson Cancer Center (P30 CA016672) and by the Molecular Genomics Core Facility and the Bioinformatics and Biostatistics Shared Resource at the H. Lee Moffitt Cancer Center & Research Institute, an NCI designated Comprehensive Cancer Center (P30-CA076292).

## Disclosure of Conflicts of Interest

**NSS:** Has intellectual property rights in the field of cellular immunotherapy and microbiome. **SSN:** Received personal fees from Kite, a Gilead Company, Merck, Bristol Myers Squibb, Novartis, Celgene, Pfizer, Allogene Therapeutics, Cell Medica/Kuur, Incyte, Precision Biosciences, Legend Biotech, Adicet Bio, Calibr, Bluebird Bio, and Unum Therapeutics; research support from Kite, a Gilead Company, Bristol Myers Squibb, Merck, Poseida, Cellectis, Celgene, Karus Therapeutics, Unum Therapeutics, Allogene Therapeutics, Precision Biosciences, and Acerta; royalties from Takeda Pharmaceuticals; and has intellectual property rights related to cell therapy. **DAS:** Received research funding from Aprea and Jazz, done consulting for AbbVie, Agios, Aprea, BMS, Incyte, Intellia, Kite, Magenta, Novartis, Shattuck Labs and Takeda and part of a speaker’s bureau for BMS, Incyte. **EP:** Obtains research funding from Incyte, Kura Oncology and BMS and has received Honoraria from Taiho. **FLL:** Scientific Advisory Role: Allogene, Amgen, Bluebird Bio, BMS/Celgene, Calibr, Cellular Biomedicine Group, GammaDelta Therapeutics, Iovance, Kite Pharma, Janssen, Legend Biotech, Novartis, Takeda, Wugen, Umoja; Research Funding: Kite Pharma (Institutional), Allogene (Institutional), Novartis (Institutional), BlueBird Bio (Institutional); Patents, Royalties, Other Intellectual Property: Several patents held by the institution in my name (unlicensed) in the field of cellular immunotherapy. **MRG** reports research funding from Sanofi, Kite/Gilead, Abbvie and Allogene, honoraria from Tessa Therapeutics and Daiichi Sankyo, and stock ownership of KDAc Therapeutics. **PS:** research support from Astrazeneca-Acerta and from ALX Oncology and consultant for Roche-Genentech and Hutchinson MediPharma. **KT**: consulting for Symbio Pharmaceuticals, Novartis, GSK, and Celgene/BMS.

## Notes

### Competing Interest Statement

The authors have declared no competing interest.

