## Supplementary File 1 for "Clonal hematopoiesis is associated with increased toxicity in large B-cell lymphoma patients treated with chimeric antigen receptor T cell therapy"

**Saini et. al.**

### SUPPLEMENTARY METHODS

#### Clonal hematopoietic (CH) detection in the MD Anderson Cancer Center cohort

For the MDACC cohort, the pre-treatment buffy coat samples were sequenced using a SureSelect custom panel of 300 genes (Agilent Technologies, Santa Clara, CA) that covers genes recurrently mutated in CH and hematologic malignancies[1]. Details of the sequencing methods have been described previously[1]. The complete descriptions of the statistical method and bioinformatics pipelines identifying high-confidence somatic single-nucleotide variants and indels from targeted capture DNA sequencing are provided below:

#### Bioinformatic algorithms to detect CHIP variants from sequenced data

Raw sequencing data from the Illumina platform were converted to a fastq format and aligned to the reference genome (hg19) using the Burroughs-Wheeler Aligner (BWA). BWA is using MEM mode with following parameters: -k 31 -T 100 -t 8 -M. The aligned BAM files were subjected to mark duplication, re-alignment, and re-calibration using Picard and GATK with default parameters (https://www.broadinstitute.org/gatk/guide/best-practices?bpm=DNAseq). Preprocessed BAM files were then analyzed to detect single nucleotide variants (SNV) and small insertions and deletions (indels) using MuTect (https://pubmed.ncbi.nlm.nih.gov/19451168/) and Pindel (https://pubmed.ncbi.nlm.nih.gov/19561018/) algorithms, respectively, against unmatched pooled normal samples.

We performed a series of filtering and annotations to identify high-confidence CHIP mutations. First, variants matching one or more of the following criteria were considered of low quality and therefore filtered out from further analysis: 1) tumor coverage < 15X, 2) tumor allele frequency < 2%, and 3) normal allele frequency ≥ 1% and 0% for SNVs and INDELs, respectively. Second, only variants which would introduce an obvious protein-coding change were kept for further analysis. Specifically, variants with an ANNOVAR annotation of non-synonymous, stop-gain, stop-loss, splicing, frameshift insertion, frameshift deletion, nonframeshift insertion or nonframeshift deletion were considered to be able to introduce an obvious protein-coding change and were therefore kept for further analysis. Third, common polymorphisms were removed to reduce the load of possible germline contamination due to the absence of matched normal. Specifically, a series of public variant databases including the 1000 Genome Database (http://www.1000genomes.org/), ESP6500 Database (http://evs.gs.washington.edu/EVS/), dbSNP ver.129 (http://www.ncbi.nlm.nih.gov/SNP/), and Exome Aggregation Consortium database (http://exac.broadinstitute.org/), were utilized. Variants with a population frequency of 0.14% or more in any of the databases were considered possible germline polymorphisms and were therefore removed from further analysis. Next, a hierarchical classification system was developed to assign a confidence level for each remaining variant in order to facilitate the identification of putative driver mutations. Specifically, each variant was classified based on the following hierarchical order and was assigned a confidence level corresponding to its rank in the system: 1) Confirmed somatic mutation based on COSMIC database (version 81), 2) loss-of-function mutation such as splicing, stop-gain, stop-loss and frameshift mutation in tumor suppressor genes, 3) variant which resides in the same position as a confirmed somatic mutation according to the COSMIC database, 4) recurrent variant which resides within three amino acids away from a confirmed somatic mutation according to the COSMIC database and was predicted to be damaging by in-silico function prediction algorithms. Finally, we used the variant filtering criteria previously used for CHIP detection, in order to identify CHIP variants [2,3]. The final annotated variant list was then further analyzed by manual inspection in order to identify the CHIP mutation.

Clonal hematopoiesis detection in Moffitt samples

DNA was extracted from peripheral blood for library preparation using our custom 76-gene hybrid-capture panel with unique molecular barcodes. This panel, combined with deep next-generation sequencing (NGS), optimally captures whole exons of all commonly mutated CH genes, including single nucleotide mutations and small indels down to allele frequencies ≥ 1%[4]. Library preparation and sequencing were conducted in collaboration with the Moffitt Molecular Genomics Core Facility. DNA libraries were generated using the SureSelect$^{XTHS}$ kit (Agilent, Santa Clara, CA) and sequenced on a NextSeq 2000 sequencer (Illumina, San Diego, CA) per manufactures' recommendations, with a goal of achieving an average coverage > 1000x. CH mutations were identified following our previously published bioinformatics pipeline[4,5]. Briefly, sequencing reads were aligned to the human genome (GRCh38) using the BWA-MEM algorithm[6]. Somatic variant calling was performed using Genome Analysis Toolkit best practices[7]. Variants were filtered and annotated using BCFtools[8]. Mutations that occurred in greater than one-third of the samples were considered sequencing artifacts and removed. Additional quality control filters included read depth > 50, strand odds ratio < 3, TLOD > 6.3, observation of each variant more than once on the forward and reverse reads, and masking of the repetitive regions of the genome as defined by the DUST algorithm[9]. Germline variants were removed using publicly available reference populations (i.e., variants observed in non-cancer populations at a frequency > 0.005 and with a VAF > 0.45 were removed)[10]. To filter for likely functional somatic variants, only mutations or indels located in exonic regions were considered; synonymous mutations and nonsynonymous mutations that had not been annotated in the Catalogue of Somatic Mutations in Cancer (COSMIC) database[11] were excluded. Remaining variants with a VAF > 0.02 were considered CH mutations.

**Table S1. The complete list of genes and variants of clonal hematopoiesis detected in 42 patients.**

| No. | gene | Mutation type | DNA/Amino Acid change | VAF |
|---|---|---|---|---|
| **1** | *ASXL1* | frameshift insertion | ASXL1:uc010geb.3:exon9:c.1600dupG:p.G533fs,ASXL1:uc002wxs.3:exon12:c.1924dupG:p.G641fs,ASXL1:uc021wbw.1:exon13:c.1927dupG:p.G642fs | 0.0292 |
| **2** | *PPM1D* | stopgain | PPM1D:uc002iyt.2:exon6:c.T1538A:p.L513X | 0.1053 |
| **3** | *PPM1D* | frameshift insertion | PPM1D:uc002iyt.2:exon6:c.1512_1513insT:p.T504fs | 0.0229 |
| **4** | *DNMT3A* | stopgain | DNMT3A:uc002rgb.4:exon5:c.C510G:p.Y170X,DNMT3A:uc002rgc.4:exon9:c.C1077G:p.Y359X,DNMT3A:uc002rgd.4:exon9:c.C1077G:p.Y359X | 0.0765 |
| **5** | *PPM1D* | stopgain | PPM1D:uc002iyt.2:exon6:c.G1270T:p.E424X | 0.0422 |
| **6** | *PPM1D* | stopgain | PPM1D:uc002iyt.2:exon6:c.C1654T:p.R552X | 0.1151 |
| **7** | *TP53* | nonsynonymous SNV | TP53:uc002gii.2:exon3:c.T218C:p.I73T,TP53:uc010cnf.2:exon3:c.T218C:p.I73T,TP53:uc010cng.2:exon3:c.T218C:p.I73T,TP53:uc002gio.3:exon4:c.T299C:p.I100T,TP53:uc002gig.1:exon6:c.T695C:p.I232T,TP53:uc002gih.3:exon6:c.T695C:p.I232T,TP53:uc002gin.3:exon6:c.T416C:p.I139T,TP53:uc031qyq.1:exon6:c.T578C:p.I193T,TP53:uc002gij.3:exon7:c.T578C:p.I193T,TP53:uc002gim.3:exon7:c.T695C:p.I232T,TP53:uc010cnh.2:exon7:c.T695C:p.I232T,TP53:uc010cni.2:exon7:c.T695C:p.I232T | 0.0798 |
| **8** | *TP53* | nonsynonymous SNV | TP53:uc002gii.2:exon3:c.G266A:p.R89Q,TP53:uc010cnf.2:exon3:c.G266A:p.R89Q,TP53:uc010cng.2:exon3:c.G266A:p.R89Q,TP53:uc002gio.3:exon4:c.G347A:p.R116Q,TP53:uc002gig.1:exon6:c.G743A:p.R248Q,TP53:uc002gih.3:exon6:c.G743A:p.R248Q,TP53:uc002gin.3:exon6:c.G464A:p.R155Q,TP53:uc031qyq.1:exon6:c.G626A:p.R209Q,TP53:uc002gij.3:exon7:c.G626A:p.R209Q,TP53:uc002gim.3:exon7:c.G743A:p.R248Q,TP53:uc010cnh.2:exon7:c.G743A:p.R248Q,TP53:uc010cni.2:exon7:c.G743A:p.R248Q | 0.0723 |
| **9** | *DNMT3A* | frameshift deletion | DNMT3A:uc002rgb.4:exon12:c.1328delA:p.K443fs,DNMT3A:uc002rgc.4:exon16:c.1895delA:p.K632fs,DNMT3A:uc002rgd.4:exon16:c.1895delA:p.K632fs | 0.4171 |
| **10** | *TP53* | nonsynonymous SNV | TP53:uc002gii.2:exon4:c.G341A:p.R114H,TP53:uc010cnf.2:exon4:c.G341A:p.R114H,TP53:uc010cng.2:exon4:c.G341A:p.R114H,TP53:uc002gih.3:exon7:c.G818A:p.R273H,TP53:uc031qyq.1:exon7:c.G701A:p.R234H,TP53:uc002gij.3:exon8:c.G701A:p.R234H,TP53:uc002gim.3:exon8:c.G818A:p.R273H,TP53:uc010cnh.2:exon8:c.G818A:p.R273H,TP53:uc010cni.2:exon8:c.G818A:p.R273H | 0.3731 |
| **11** | *TP53* | nonsynonymous SNV | TP53:uc002gii.2:exon6:c.T554C:p.L185P,TP53:uc031qyq.1:exon9:c.T914C:p.L305P,TP53:uc002gij.3:exon10:c.T914C:p.L305P,TP53:uc002gim.3:exon10:c.T1031C:p.L344P | 0.2836 |
| **12** | *PPM1D* | stopgain | PPM1D:uc002iyt.2:exon6:c.C1434A:p.C478X | 0.0277 |
| **13** | *TP53* | nonsynonymous SNV | TP53:uc002gii.2:exon4:c.C340T:p.R114C,TP53:uc010cnf.2:exon4:c.C340T:p.R114C,TP53:uc010cng.2:exon4:c.C340T:p.R114C,TP53:uc002gih.3:exon7:c.C817T:p.R273C,TP53:uc031qyq.1:exon7:c.C700T:p.R234C,TP53:uc002gij.3:exon8:c.C700T:p.R234C,TP53:uc002gim.3:exon8:c.C817T:p.R273C,TP53:uc010cnh.2:exon8:c.C817T:p.R273C,TP53:uc010cni.2:exon8:c.C817T:p.R273C | 0.0437 |
| **14** | *TP53* | nonsynonymous SNV | TP53:uc002gii.2:exon2:c.T168G:p.S56R,TP53:uc010cnf.2:exon2:c.T168G:p.S56R,TP53:uc010cng.2:exon2:c.T168G:p.S56R,TP53:uc002gio.3:exon3:c.T249G:p.S83R,TP53:uc002gig.1:exon5:c.T645G:p.S215R,TP53:uc002gih.3:exon5:c.T645G:p.S215R,TP53:uc002gin.3:exon5:c.T366G:p.S122R,TP53:uc010vug.3:exon5:c.T528G:p.S176R,TP53:uc031qyq.1:exon5:c.T528G:p.S176R,TP53:uc002gij.3:exon6:c.T528G:p.S176R,TP53:uc002gim.3:exon6:c.T645G:p.S215R,TP53:uc010cnh.2:exon6:c.T645G:p.S215R,TP53:uc010cni.2:exon6:c.T645G:p.S215R | 0.1127 |
| **15** | *PPM1D* | frameshift deletion | PPM1D:uc002iyt.2:exon6:c.1529delA:p.Q510fs | 0.021 |
| **16** | *ASXL1* | frameshift insertion | ASXL1:uc010geb.3:exon9:c.1600dupG:p.G533fs,ASXL1:uc002wxs.3:exon12:c.1924dupG:p.G641fs,ASXL1:uc021wbw.1:exon13:c.1927dupG:p.G642fs | 0.1095 |
| **17** | *PPM1D* | frameshift insertion | PPM1D:uc002iyt.2:exon6:c.1450_1451insTA:p.L484fs | 0.3333 |
| **18** | *TP53* | nonsynonymous SNV | TP53:uc002gio.3:exon2:c.G73T:p.V25F,TP53:uc002gig.1:exon4:c.G469T:p.V157F,TP53:uc002gih.3:exon4:c.G469T:p.V157F,TP53:uc002gin.3:exon4:c.G190T:p.V64F,TP53:uc010vug.3:exon4:c.G352T:p.V118F,TP53:uc031qyq.1:exon4:c.G352T:p.V118F,TP53:uc002gij.3:exon5:c.G352T:p.V118F,TP53:uc002gim.3:exon5:c.G469T:p.V157F,TP53:uc010cnh.2:exon5:c.G469T:p.V157F,TP53:uc010cni.2:exon5:c.G469T:p.V157F | 0.0579 |
| **19** | *TET2* | stopgain | TET2:uc003hxk.3:exon11:c.T4983G:p.Y1661X,TET2:uc011cez.2:exon11:c.T5046G:p.Y1682X | 0.0392 |
| **20** | *PPM1D* | frameshift deletion | PPM1D:uc002iyt.2:exon6:c.1529delA:p.Q510fs | 0.08 |
| **21** | *PPM1D* | frameshift insertion | PPM1D:uc002iyt.2:exon6:c.1710dupA:p.S570fs | 0.0618 |
| **22** | *PPM1D* | frameshift deletion | PPM1D:uc002iyt.2:exon6:c.1433delG:p.C478fs | 0.0205 |

| | | | | |
|---|---|---|---|---|
| **23** | *PPM1D* | frameshift insertion | PPM1D:uc002iyt.2:exon6:c.1529dupA:p.Q510fs | 0.0479 |
| **24** | *PPM1D* | stopgain | PPM1D:uc002iyt.2:exon6:c.C1741T:p.R581X | 0.0320 |
| **25** | *TP53* | nonsynonymous SNV | TP53:uc002gii.2:exon4:c.G341A:p.R114H,TP53:uc010cnf.2:exon4:c.G341A:p.R114H,TP53:uc010cng.2:exon4:c.G341A:p.R114H,TP53:uc002gih.3:exon7:c.G818A:p.R273H,TP53:uc031qyq.1:exon7:c.G701A:p.R234H,TP53:uc002gij.3:exon8:c.G701A:p.R234H,TP53:uc002gim.3:exon8:c.G818A:p.R273H,TP53:uc010cnh.2:exon8:c.G818A:p.R273H,TP53:uc010cni.2:exon8:c.G818A:p.R273H | 0.2 |
| **26** | *TET2* | frameshift deletion | TET2:uc003hxk.3:exon11:c.4984delC:p.P1662fs,TET2:uc011cez.2:exon11:c.5047delC:p.P1683fs | 0.0406 |
| **27** | *TP53* | nonsynonymous SNV | TP53:uc002gii.2:exon1:c.A59T:p.H20L,TP53:uc010cnf.2:exon1:c.A59T:p.H20L,TP53:uc010cng.2:exon1:c.A59T:p.H20L,TP53:uc002gio.3:exon2:c.A140T:p.H47L,TP53:uc002gig.1:exon4:c.A536T:p.H179L,TP53:uc002gih.3:exon4:c.A536T:p.H179L,TP53:uc002gin.3:exon4:c.A257T:p.H86L,TP53:uc010vug.3:exon4:c.A419T:p.H140L,TP53:uc031qyq.1:exon4:c.A419T:p.H140L,TP53:uc002gij.3:exon5:c.A419T:p.H140L,TP53:uc002gim.3:exon5:c.A536T:p.H179L,TP53:uc010cnh.2:exon5:c.A536T:p.H179L,TP53:uc010cni.2:exon5:c.A536T:p.H179L | 0.0385 |
| **28** | *PPM1D* | frameshift insertion | PPM1D:uc002iyt.2:exon6:c.1449dupT:p.T483fs | 0.0764 |
| **29** | *TP53* | nonsynonymous SNV | TP53:uc002gii.2:exon3:c.A238G:p.N80D,TP53:uc010cnf.2:exon3:c.A238G:p.N80D,TP53:uc010cng.2:exon3:c.A238G:p.N80D,TP53:uc002gio.3:exon4:c.A319G:p.N107D,TP53:uc002gig.1:exon6:c.A715G:p.N239D,TP53:uc002gih.3:exon6:c.A715G:p.N239D,TP53:uc002gin.3:exon6:c.A436G:p.N146D,TP53:uc031qyq.1:exon6:c.A598G:p.N200D,TP53:uc002gij.3:exon7:c.A598G:p.N200D,TP53:uc002gim.3:exon7:c.A715G:p.N239D,TP53:uc010cnh.2:exon7:c.A715G:p.N239D,TP53:uc010cni.2:exon7:c.A715G:p.N239D | 0.0476 |
| **30** | *PPM1D* | frameshift deletion | PPM1D:uc002iyt.2:exon6:c.1286delG:p.R429fs | 0.0345 |
| **31** | *PPM1D* | frameshift deletion | PPM1D:uc002iyt.2:exon6:c.1632delC:p.G544fs | 0.0333 |
| **32** | *PPM1D* | frameshift deletion | PPM1D:uc002iyt.2:exon6:c.1529delA:p.Q510fs | 0.0702 |
| **33** | *ASXL1* | frameshift insertion | ASXL1:uc010geb.3:exon9:c.1600dupG:p.G533fs,ASXL1:uc002wxs.3:exon12:c.1924dupG:p.G641fs,ASXL1:uc021wbw.1:exon13:c.1927dupG:p.G642fs | 0.0345 |
| **34** | *ASXL1* | nonframeshift deletion | ASXL1:uc010geb.3:exon9:c.1581_1604del:p.527_535del,ASXL1:uc002wxs.3:exon12:c.1905_1928del:p.635_643del,ASXL1:uc021wbw.1:exon13:c.1908_1931del:p.636_644del | 0.0519 |
| **35** | *TP53* | frameshift deletion | TP53:uc002gii.2:exon6:c.520delC:p.R174fs,TP53:uc031qyq.1:exon9:c.880delC:p.R294fs,TP53:uc002gij.3:exon10:c.880delC:p.R294fs,TP53:uc002gim.3:exon10:c.997delC:p.R333fs | 0.2025 |
| **36** | *PPM1D* | stopgain | PPM1D:uc002iyt.2:exon6:c.C1741T:p.R581X | 0.1259 |
| **37** | *PPM1D* | frameshift deletion | PPM1D:uc002iyt.2:exon6:c.1586delC:p.T529fs | 0.0473 |
| **38** | *PPM1D* | stopgain | PPM1D:uc002iyt.2:exon6:c.C1741T:p.R581X | 0.1818 |
| **39** | *PPM1D* | frameshift deletion | PPM1D:uc002iyt.2:exon6:c.1586delC:p.T529fs | 0.1032 |
| **40** | *TP53* | frameshift deletion | TP53:uc002gii.2:exon6:c.520delC:p.R174fs,TP53:uc031qyq.1:exon9:c.880delC:p.R294fs,TP53:uc002gij.3:exon10:c.880delC:p.R294fs,TP53:uc002gim.3:exon10:c.997delC:p.R333fs | 0.0274 |
| **41** | *TET2* | frameshift deletion | TET2:uc021xql.1:exon1:c.2909delC:p.T970fs,TET2:uc010ilp.2:exon2:c.2909delC:p.T970fs,TET2:uc003hxk.3:exon3:c.2909delC:p.T970fs,TET2:uc011cez.2:exon3:c.2972delC:p.T991fs,TET2:uc021xqk.1:exon3:c.2909delC:p.T970fs | 0.1034 |
| **42** | *TET2* | frameshift deletion | TET2:uc021xql.1:exon1:c.2779delG:p.V927fs,TET2:uc010ilp.2:exon2:c.2779delG:p.V927fs,TET2:uc003hxk.3:exon3:c.2779delG:p.V927fs,TET2:uc011cez.2:exon3:c.2842delG:p.V948fs,TET2:uc021xqk.1:exon3:c.2779delG:p.V927fs | 0.0703 |
| **43** | *GNAS* | nonsynonymous SNV | GNAS:uc002yae.3:exon5:c.G377A:p.R126H,GNAS:uc002yad.3:exon6:c.G275A:p.R92H,GNAS:uc002yaa.3:exon7:c.G557A:p.R186H,GNAS:uc021wfp.1:exon7:c.G560A:p.R187H,GNAS:uc002xzt.3:exon8:c.G506A:p.R169H,GNAS:uc002xzw.3:exon8:c.G2531A:p.R844H,GNAS:uc002xzx.3:exon8:c.G425A:p.R142H,GNAS:uc010gjq.3:exon8:c.G425A:p.R142H,GNAS:uc021wfn.1:exon8:c.G602A:p.R201H,GNAS:uc021wfo.1:exon8:c.G605A:p.R202H | 0.0253 |
| **44** | *PPM1D* | frameshift insertion | PPM1D:uc002iyt.2:exon6:c.1626dupT:p.N542fs | 0.037 |
| **45** | *PPM1D* | frameshift deletion | PPM1D:uc002iyt.2:exon6:c.1437delT:p.A479fs | 0.06 |
| **46** | *TP53* | nonsynonymous SNV | TP53:uc002gii.2:exon2:c.A101G:p.H34R,TP53:uc010cnf.2:exon2:c.A101G:p.H34R,TP53:uc010cng.2:exon2:c.A101G:p.H34R,TP53:uc002gio.3:exon3:c.A182G:p.H61R,TP53:uc002gig.1:exon5:c.A578G:p.H193R,TP53:uc002gih.3:exon5:c.A578G:p.H193R,TP53:uc002gin.3:exon5:c.A299G:p.H100R,TP53:uc010vug.3:exon5:c.A461G:p.H154R,TP53:uc031qyq.1:exon5:c.A461G:p.H154R,TP53:uc002gij.3:exon6:c.A461G:p.H154R,TP53:uc002gim.3:exon6:c.A578G:p.H193R,TP53:uc010cnh.2:exon6:c.A578G:p.H193R,TP53:uc010cni.2:exon6:c.A578G:p.H193R | 0.0477 |

| | | | | |
|---|---|---|---|---|
| **47** | *IDH2* | nonsynonymous SNV | IDH2:uc010uqc.2:exon2:c.G29A:p.R10Q,IDH2:uc002box.3:exon4:c.G419A:p.R140Q,IDH2:uc010uqb.2:exon4:c.G263A:p.R88Q | 0.0245 |
| **48** | *PPM1D* | frameshift deletion | PPM1D:uc002iyt.2:exon6:c.1387delG:p.G463fs | 0.0679 |
| **49** | *PPM1D* | frameshift deletion | PPM1D:uc002iyt.2:exon6:c.1529delA:p.Q510fs | 0.0658 |
| **50** | *PPM1D* | frameshift insertion | PPM1D:uc002iyt.2:exon6:c.1433dupG:p.C478fs | 0.0443 |
| **51** | *PPM1D* | stopgain | PPM1D:uc002iyt.2:exon6:c.C1358A:p.S453X | 0.0408 |
| **52** | *TP53* | nonsynonymous SNV | TP53:uc002gii.2:exon3:c.G256A:p.G86S,TP53:uc010cnf.2:exon3:c.G256A:p.G86S,TP53:uc010cng.2:exon3:c.G256A:p.G86S,TP53:uc002gio.3:exon4:c.G337A:p.G113S,TP53:uc002gig.1:exon6:c.G733A:p.G245S,TP53:uc002gih.3:exon6:c.G733A:p.G245S,TP53:uc002gin.3:exon6:c.G454A:p.G152S,TP53:uc031qyq.1:exon6:c.G616A:p.G206S,TP53:uc002gij.3:exon7:c.G616A:p.G206S,TP53:uc002gim.3:exon7:c.G733A:p.G245S,TP53:uc010cnh.2:exon7:c.G733A:p.G245S,TP53:uc010cni.2:exon7:c.G733A:p.G245S | 0.0790 |
| **53** | *PPM1D* | frameshift insertion | PPM1D:uc002iyt.2:exon6:c.1543_1544insTG:p.M515fs | 0.2978 |
| **54** | *DNMT3A* | nonsynonymous SNV | DNMT3A:uc002rgb.4:exon19:c.C2077T:p.R693C,DNMT3A:uc002rgc.4:exon23:c.C2644T:p.R882C,DNMT3A:uc002rgd.4:exon23:c.C2644T:p.R882C | 0.0475 |
| **55** | *TP53* | nonsynonymous SNV | TP53:uc002gii.2:exon2:c.T168G:p.S56R,TP53:uc010cnf.2:exon2:c.T168G:p.S56R,TP53:uc010cng.2:exon2:c.T168G:p.S56R,TP53:uc002gio.3:exon3:c.T249G:p.S83R,TP53:uc002gig.1:exon5:c.T645G:p.S215R,TP53:uc002gih.3:exon5:c.T645G:p.S215R,TP53:uc002gin.3:exon5:c.T366G:p.S122R,TP53:uc010vug.3:exon5:c.T528G:p.S176R,TP53:uc031qyq.1:exon5:c.T528G:p.S176R,TP53:uc002gij.3:exon6:c.T528G:p.S176R,TP53:uc002gim.3:exon6:c.T645G:p.S215R,TP53:uc010cnh.2:exon6:c.T645G:p.S215R,TP53:uc010cni.2:exon6:c.T645G:p.S215R | 0.0558 |
| **56** | *TET2* | frameshift deletion | TET2:uc021xql.1:exon1:c.944delC:p.S315fs,TET2:uc010ilp.2:exon2:c.944delC:p.S315fs,TET2:uc003hxk.3:exon3:c.944delC:p.S315fs,TET2:uc011cez.2:exon3:c.1007delC:p.S336fs,TET2:uc021xqk.1:exon3:c.944delC:p.S315fs | 0.0372 |
| **57** | *PPM1D* | stopgain | PPM1D:uc002iyt.2:exon6:c.G1280A:p.W427X | 0.0382 |
| **58** | *BCOR* | frameshift deletion | BCOR:uc004dem.4:exon4:c.692_693del:p.Q231fs,BCOR:uc004den.4:exon4:c.692_693del:p.Q231fs,BCOR:uc004deo.4:exon4:c.692_693del:p.Q231fs,BCOR:uc004dep.4:exon4:c.692_693del:p.Q231fs,BCOR:uc004deq.4:exon4:c.692_693del:p.Q231fs | 0.2706 |
| **59** | *PPM1D* | frameshift deletion | PPM1D:uc002iyt.2:exon6:c.1333delT:p.F445fs | 0.163 |
| **60** | *PPM1D* | stopgain | PPM1D:uc002iyt.2:exon6:c.C1741T:p.R581X | 0.0281 |
| **61** | *DNMT3A* | stopgain | DNMT3A:uc002rgb.4:exon10:c.C1012T:p.Q338X,DNMT3A:uc002rgc.4:exon14:c.C1579T:p.Q527X,DNMT3A:uc002rgd.4:exon14:c.C1579T:p.Q527X | 0.0320 |
| **62** | *U2AF1* | nonsynonymous SNV | U2AF1:uc002zda.1:exon2:c.C101A:p.S34Y,U2AF1:uc002zdb.1:exon2:c.C101A:p.S34Y,U2AF1:uc002zdc.1:exon2:c.C101A:p.S34Y,U2AF1:uc010gpi.1:exon2:c.C101A:p.S34Y | 0.2421 |
| **63** | *PPM1D* | stopgain | PPM1D:uc002iyt.2:exon6:c.C1654T:p.R552X | 0.0328 |
| **64** | *PPM1D* | frameshift deletion | PPM1D:uc002iyt.2:exon6:c.1467delT:p.S489fs | 0.0296 |
| **65** | *TET2* | frameshift deletion | TET2:uc021xql.1:exon1:c.1606delA:p.K536fs,TET2:uc010ilp.2:exon2:c.1606delA:p.K536fs,TET2:uc003hxk.3:exon3:c.1606delA:p.K536fs,TET2:uc011cez.2:exon3:c.1669delA:p.K557fs,TET2:uc021xqk.1:exon3:c.1606delA:p.K536fs | 0.0825 |
| **66** | *PPM1D* | frameshift insertion | PPM1D:uc002iyt.2:exon6:c.1529dupA:p.Q510fs | 0.0765 |
| **67** | *PPM1D* | frameshift insertion | PPM1D:uc002iyt.2:exon6:c.1449dupT:p.T483fs | 0.0526 |
| **68** | *DNMT3A* | frameshift deletion | DNMT3A:uc002rgb.4:exon5:c.488delG:p.S163fs,DNMT3A:uc002rgc.4:exon9:c.1055delG:p.S352fs,DNMT3A:uc002rgd.4:exon9:c.1055delG:p.S352fs | 0.05 |
| **69** | *TET2* | frameshift deletion | TET2:uc003hxk.3:exon11:c.4851delT:p.P1617fs,TET2:uc011cez.2:exon11:c.4914delT:p.P1638fs | 0.0677 |
| **70** | *CHEK2* | stopgain | CHEK2:uc010gvi.1:exon3:c.467dupA:p.Y156_I157delinsX,CHEK2:uc003adu.1:exon4:c.467dupA:p.Y156_I157delinsX,CHEK2:uc003adv.1:exon4:c.467dupA:p.Y156_I157delinsX,CHEK2:uc003adt.1:exon5:c.596dupA:p.Y199_I200delinsX | 0.0613 |
| **71** | *DNMT3A* | nonsynonymous SNV | p.S770L | 0.215 |
| **72** | *DNMT3A* | nonsynonymous SNV | p.S770L | 0.213 |

**Table S2. Univariate Cox analysis for progression-free survival with CH mutations**

| Variables | Hazard ratio | P-value |
|---|---|---|
| CH mutations absent | 0.97 (0.62 - 1.53) | 0.901 |
| *DNMT3A* | 1.26 (0.53 - 3.0) | 0.605 |
| *PPM1D* | 1.05 (0.58 – 1.91) | 0.877 |
| *TP53* | 1.27 (0.66 – 2.46) | 0.470 |
| *TET2* | 0.85 (0.31 – 2.38) | 0.763 |

**Table S3. Univariate Cox analysis for overall survival with CH mutations**

| Variables | Hazard ratio | P-value |
|---|---|---|
| CH mutations absent | 0.75(0.44- 1.28) | 0.295 |
| *DNMT3A* | 1.75 (0.68 – 4.53) | 0.316 |
| *PPM1D* | 0.85 (0.31 – 2.38) | 0.763 |
| *TP53* | 1.23 (0.54 – 2.79) | 0.621 |
| *TET2* | 1.20 (0.36 – 3.93) | 0.767 |

**Table S4. Univariate analysis for covariates of interest with severe grade ICANS**

| Variable | N | Overall | ICANS ≤2 (n=77) | ICANS ≥3 (n=37) | P |
|---|---|---|---|---|---|
| n (%) | 114 | | | | 0.038 |
| CH | | 42 (36.8) | 23 (29.9) | 19 (51.4) | |
| No CH | | 72 (63.2) | 54 (70.1) | 18 (48.6) | |
| Age, M (SD) | 114 | 60.5 (12.9) | 60.2 (13.0) | 61.2 (12.8) | 0.701 |
| Sex, n (%) | 114 | | | | 0.392 |
| Female | | 34 (29.8) | 21 (27.3) | 13 (35.1) | |
| Male | | 80 (70.2) | 56 (72.7) | 24 (64.9) | |
| Histo, n (%) | 114 | | | | 1.000 |
| DLBCL | | 91 (79.8) | 61 (79.2) | 30 (81.1) | |
| TFL/PMBCL | | 23 (20.2) | 16 (20.8) | 7 (18.9) | |
| ECOG, n (%) | 113 | | | | 0.668 |
| =0 | | 34 (30.1) | 24 (31.6) | 10 (27.0) | |
| >0 | | 79 (69.9) | 52 (68.4) | 27 (73.0) | |
| Stage, n (%) | 114 | | | | 0.797 |
| I/II | | 20 (17.5) | 13 (16.9) | 7 (18.9) | |
| III/IV | | 94 (82.5) | 64 (83.1) | 30 (81.1) | |
| IPIscore, n (%) | 114 | | | | 0.683 |
| <=2 | | 44 (38.6) | 31 (40.3) | 13 (35.1) | |
| 3-5 | | 70 (61.4) | 46 (59.7) | 24 (64.9) | |
| Ferritin, M (SD) | 106 | 1719.8 (4167.7) | 1022.6 (1486.7) | 3075.5 (6699.9) | 0.078 |
| LDH, M (SD) | 106 | 355.8 (370.8) | 304.5 (139.9) | 455.8 (598.6) | 0.143 |
| CRP, M (SD) | 105 | 45.0 (62.5) | 35.5 (55.8) | 63.0 (71.0) | 0.048 |
| EGFR, n (%) | 105 | | | | 0.199 |
| <=60 | | 21 (20.0) | 11 (15.9) | 10 (27.8) | |
| >60 | | 84 (80.0) | 58 (84.1) | 26 (72.2) | |
| Line.Prevs, M (SD) | 114 | 3.7 (1.7) | 3.7 (1.9) | 3.7 (1.4) | 0.930 |
| Refractory, n (%) | 114 | | | | 0.243 |
| No | | 27 (23.7) | 21 (27.3) | 6 (16.2) | |
| Yes | | 87 (76.3) | 56 (72.7) | 31 (83.8) | |
| ASCT.prevs, n (%) | 114 | | | | 1.000 |
| No | | 89 (78.1) | 60 (77.9) | 29 (78.4) | |
| Yes | | 25 (21.9) | 17 (22.1) | 8 (21.6) | |

**Table S5. Multivariate analysis for covariates of interest with severe grade ICANS**

| Variable | OR | 95% CI for OR | P |
|---|---|---|---|
| CH<br>No CH (ref) | - | - | - |
| CH | 2.47 | (1.02, 6.02) | 0.046 |
| Age | 1.00 | (0.96, 1.04) | 0.876 |
| Ferritin | 1.00 | (1.00, 1.00) | 0.116 |
| LDH | 1.00 | (1.00, 1.00) | 0.842 |
| CRP | 1.00 | (1.00, 1.01) | 0.386 |
| EGFR<br><=60 (ref) | - | - | - |
| >60 | 0.58 | (0.19, 1.78) | 0.343 |
| Refractory<br>No (ref) | - | - | - |
| Yes | 1.12 | (0.37, 3.41) | 0.839 |

**Table S6: Serum cytokines measurement at DAY 0 in available samples in CH and no-CH patients.**

| Variable | N | Overall | CH (n=14) | No CH (n=14) | P |
|---|---|---|---|---|---|
| GM-CSF, Median (Min-Max) | 27 | 0.07 (0.00-1.64) | 0.07 (0.00-0.85) | 0.08 (0.00-1.64) | 0.884 |
| IFN-γ, Median (Min-Max) | 28 | 3.43 (0.00-123.53) | 3.14 (0.00-79.50) | 4.75 (0.00-123.53) | 0.268 |
| IL-10, Median (Min-Max) | 28 | 0.27 (0.00-10.69) | 0.27 (0.00-10.69) | 0.26 (0.00-4.98) | 0.730 |
| IL-13, Median (Min-Max) | 28 | 0.05 (0.00-32.68) | 0.23 (0.00-14.19) | 0.00 (0.00-32.68) | 0.445 |
| IL-17A, Median (Min-Max) | 28 | 0.67 (0.00-24.61) | 0.73 (0.00-10.60) | 0.46 (0.00-24.61) | 0.230 |
| IL-2, Median (Min-Max) | 28 | 0.46 (0.00-45.92) | 0.63 (0.00-5.82) | 0.29 (0.00-45.92) | 0.311 |
| IL-4, Median (Min-Max) | 28 | 0.01 (0.00-0.86) | 0.02 (0.00-0.17) | 0.01 (0.00-0.86) | 0.835 |
| IL-5, Median (Min-Max) | 28 | 0.31 (0.00-30.27) | 0.27 (0.00-3.79) | 0.31 (0.00-30.27) | 0.927 |
| IL-6, Median (Min-Max) | 28 | 0.91 (0.22-8.38) | 1.12 (0.42-4.34) | 0.62 (0.22-8.38) | 0.058 |
| TNF-α, Median (Min-Max) | 28 | 0.83 (0.00-35.48) | 0.87 (0.00-6.49) | 0.75 (0.00-35.48) | 0.818 |
| IL-7, Median (Min-Max) | 24 | 14.32 (0.36-58.55) | 11.40 (0.36-35.65) | 15.10 (0.54-58.55) | 0.443 |
| IL-15, Median (Min-Max) | 24 | 18.76 (1.85-47.24) | 18.35 (1.85-24.70) | 21.15 (6.70-47.24) | 0.266 |

**Table S7: Median of the highest levels of serum cytokines post-CAR-T infusion in the two cohorts.**

| Variable | N | Overall | CH (n=19) | No CH (n=25) | P |
|---|---|---|---|---|---|
| **GM-CSF, Median (Min-Max)** | 44 | 0.93 (0.00-14.15) | 1.10 (0.10-14.15) | 0.90 (0.00-2.86) | 0.438 |
| **IFN-γ, Median (Min-Max)** | 44 | 139.12 (8.03-2858.40) | 143.37 (16.48-2536.03) | 134.09 (8.03-2858.40) | 0.778 |
| **IL-10, Median (Min-Max)** | 44 | 8.14 (0.39-308.04) | 4.01 (1.02-145.63) | 10.08 (0.39-308.04) | 0.157 |
| **IL-13, Median (Min-Max)** | 44 | 13.98 (0.00-179.24) | 13.71 (0.23-179.24) | 14.21 (0.00-66.26) | 0.831 |
| **IL-17, Median (Min-Max)** | 44 | 8.00 (0.27-123.23) | 16.43 (0.63-123.23) | 7.32 (0.27-56.89) | 0.606 |
| **IL-2, Median (Min-Max)** | 44 | 5.74 (0.76-45.92) | 4.86 (0.76-37.85) | 6.30 (1.50-45.92) | 0.411 |
| **IL-4, Median (Min-Max)** | 44 | 0.31 (0.01-4.67) | 0.21 (0.02-4.67) | 0.35 (0.01-3.06) | 0.496 |
| **IL-5, Median (Min-Max)** | 44 | 7.45 (0.62-187.79) | 11.35 (0.78-187.79) | 6.30 (0.62-37.37) | 0.279 |
| **TNF-α, Median (Min-Max)** | 44 | 6.02 (0.34-68.72) | 4.35 (0.34-68.72) | 6.43 (0.80-35.48) | 0.195 |
| **IL-6, Median (Min-Max)** | 44 | 11.19 (1.03-1825.53) | 13.27 (1.03-1825.53) | 10.96 (1.85-376.93) | 0.690 |

**Table S8. Patient blood parameter characteristics between the CH and no CH cohorts.**

| Variable | N | Overall | CH | No-CH | P |
|---|---|---|---|---|---|
| **ANC. baseline, Median (Min-Max)** | 80 | 2.8 (0.0-23.0) | 3.1 (0.9-23.0) | 2.5 (0.0-19.1) | 0.070 |
| **ALC. baseline, Median (Min-Max)** | 80 | 0.6 (0.1-2.8) | 0.5 (0.1-2.8) | 0.6 (0.1-2.4) | 0.182 |
| **AMC. baseline, Median (Min-Max)** | 80 | 0.5 (0.0-4.2) | 0.5 (0.0-4.2) | 0.5 (0.0-2.0) | 0.942 |
| **HgB. baseline, Median (Min-Max)** | 80 | 10.4 (7.3-16.5) | 10.5 (7.3-16.5) | 10.1 (7.3-14.2) | 0.622 |
| **PLT. baseline, Median (Min-Max)** | 80 | 117.0 (9.0-494.0) | 106.5 (38.0-292.0) | 119.0 (9.0-494.0) | 0.445 |
| **ANC.Day90, Median (Min-Max)** | 78 | 1.9 (0.0-9.7) | 1.5 (0.0-9.7) | 2.4 (0.0-7.6) | 0.093 |
| **ALC.Day90, Median (Min-Max)** | 76 | 0.5 (0.1-8.8) | 0.6 (0.2-2.2) | 0.5 (0.1-8.8) | 0.912 |
| **WBC.Day90, Median (Min-Max)** | 78 | 3.1 (0.8-11.2) | 3.0 (0.8-10.4) | 3.6 (1.0-11.2) | 0.119 |
| **HgB.Day90, Median (Min-Max)** | 78 | 10.4 (1.7-16.7) | 10.6 (7.7-16.7) | 10.3 (1.7-14.8) | 0.340 |
| **PLT.Day90, Median (Min-Max)** | 78 | 92.5 (3.0-313.0) | 85.0 (8.0-216.0) | 102.0 (3.0-313.0) | 0.343 |

Abbreviations: ALC – absolute lymphocyte count; AMC – absolute monocyte count; ANC -absolute neutrophil count; HgB –hemoglobin; PLT - platelets; WBC - whole blood counts

**Table S9. Clinical course and CHIP mutations detected in patients who developed MDS post CAR-T therapy infusion.**

| Patient No. | Gene | Type of mutation | Mutation region |
|---|---|---|---|
| **1** | *PPM1D* | Frameshift insertion | PPM1D:uc002iyt.2:exon6:c.1529dupA:p.Q510fs |
| | *PPM1D* | Frameshift deletion | PPM1D:uc002iyt.2:exon6:c.1449dupT:p.T483fs |
| | *TET2* | Frameshift deletion | TET2:uc021xql.1:exon1:c.1606delA:p.K536fs,TET2:uc010ilp.2:exon2:c.1606delA:p.K536fs,TET2:uc003hxk.3:exon3:c.1606delA:p.K536fs,TET2:uc011cez.2:exon3:c.1669delA:p.K557fs,TET2:uc021xqk.1:exon3:c.1606delA:p.K536fs |
| **57-year-old male with relapsed-refractory high-grade DLBCL who was in remission 8 months post CAR-T therapy – presented with pancytopenia and BM biopsy that was consistent with therapy-related MDS/MPN with 7% blasts. Leukemia mutation panel assay detected mutation on *TET2* and *TP53* gene in bone marrow. Patient was started on a clinical trial of magrolimumab, but unfortunately died a month later from respiratory distress secondary to pneumonia. PET-CT scan a month before his death showed him to be in remission from lymphoma.** | | | |
| **2** | *DNMT3A* | Frameshift deletion | DNMT3A:uc002rgb.4:exon12:c.1328delA:p.K443fs,DNMT3A:uc002rgc.4:exon16:c.1895delA:p.K632fs,DNMT3A:uc002rgd.4:exon16:c.1895delA:p.K632fs |
| **63-year-old male with relapsed refractory DLBCL was in remission for more than 2-years post CAR-T therapy. He presented in the clinic with elevated WBC up to 71K and was diagnosed with acute myeloid leukemia with 46% blasts on bone marrow biopsy. Leukemia mutation panel assay detected a mutation in *DNMT3A* (exon16:c.1895delA:p.K632fs), along with other mutations in *NRAS, KRAS, BCOR, CBL,* and *PTPN11* genes in bone marrow. He was started on a clinical trial with azacytidine plus venetoclax plus pevonidostat and unfortunately, progressed after three cycles. The patient received two more lines of therapy before succumbing to his leukemia at 30 months post CAR-T therapy.** | | | |
| **3** | *TP53* | Nonsynonymous | TP53:uc002gii.2:exon1:c.A59T:p.H20L,TP53:uc010cnf.2:exon1:c.A59T:p.H20L,TP53:uc010cng.2:exon1:c.A59T:p.H20L,TP53:uc002gio.3:exon2:c.A140T:p.H47L,TP53:uc002gig.1:exon4:c.A536T:p.H179L,TP53:uc002gih.3:exon4:c.A536T:p.H179L,TP53:uc002gin.3:exon4:c.A257T:p.H86L,TP53:uc010vug.3:exon4:c.A419T:p.H140L,TP53:uc031qyq.1:exon4:c.A419T:p.H140L,TP53:uc002gij.3:exon5:c.A419T:p.H140L,TP53:uc002gim.3:exon5:c.A536T:p.H179L |
| **87-year-old male with relapsed refractory DLBCL at 20 months remission post CAR-T therapy presented with progressive pancytopenia and transfusion dependence. His bone marrow biopsy showed a treatment related myeloid neoplasm, morphologically consistent with chronic myelomonocytic leukemia -1 with 6% blasts. Leukemia mutation panel assay at MDACC showed mutation in *TP53* (exon5:c.A536T:p.H179L, VAF<5%), along with new mutations in *SETBP1* and *ETV6* in bone marrow. He was started on azacytidine and remains alive with stable disease at 24 months post CAR-T therapy.** | | | |
| **4** | *PPM1D* | Stopgain | PPM1D:uc002iyt.2:exon6:c.C1654T:p.R552X |
| | *TP53* | Nonsynonymous SNV | TP53:uc002gii.2:exon3:c.T218C:p.I73T,TP53:uc010cnf.2:exon3:c.T218C:p.I73T,TP53:uc010cng.2:exon3:c.T218C:p.I73T,TP53:uc002gio.3:exon4:c.T299C:p.I100T,TP53:uc002gig.1:exon6:c.T695C:p.I232T,TP53:uc002gih.3:exon6:c.T695C:p.I232T,TP53:uc002gin.3:exon6:c.T416C:p.I139T,TP53:uc031qyq.1:exon6:c.T578C:p.I193T,TP53:uc002gij.3:exon7:c.T578C:p.I193T,TP53:uc002gim.3:exon7:c.T695C:p.I232T,TP53:uc010cnh.2:exon7:c.T695C:p.I232T, TP53:uc010cni.2:exon7:c.T695C:p.I232T |
| | *TP53* | Nonsynonymous SNV | TP53:uc002gii.2:exon3:c.G266A:p.R89Q,TP53:uc010cnf.2:exon3:c.G266A:p.R89Q,TP53:uc010cng.2:exon3:c.G266A:p.R89Q,TP53:uc002gio.3:exon4:c.G347A:p.R116Q,TP53:uc002gig.1:exon6:c.G743A:p.R248Q,TP53:uc002gih.3:exon6:c.G743A:p.R248Q,TP53:uc002gin.3:exon6:c.G464A:p.R155Q,TP53:uc031qyq.1:exon6:c.G626A:p.R209Q,TP53:uc002gij.3:exon7:c.G626A:p.R209Q,TP53:uc002gim.3:exon7:c.G743A:p.R248Q,TP53:uc010cnh.2:exon7:c.G743A:p.R248Q,TP53:uc010cni.2:exon7:c.G743A:p.R248Q |
| **71-year-old male with relapsed refractory DLBCL relapsed 7 months after CAR-T therapy. Later, he received 2 lines of anti-lymphoma salvage regimens with partial control of the disease. At 11 months post CAR-T infusion, he was admitted for respiratory distress secondary to pneumonia** | | | |

| | | | |
|---|---|---|---|
| **and during his hospital stay was diagnosed with therapy related AML. He was not a candidate for further therapy due to decline in his status and passed away at 13 months post CAR-t therapy from progressive lymphoma and t-AML. Leukemia mutation panel was not done in this patient.** | | | |
| **5** | *DNMT3A* | Nonsynonymous SNV | Exon19:c.C2309T:p.S770 |
| **69-year-old female with relapsed transformed follicular lymphoma developed multiorgan failure, pancytopenia and rising ferritin post CAR-T therapy. 3 months prior to CAR-T, a bone marrow was performed that showed no evidence of dysplasia or lymphoma. Bone marrow at day 30 showed treatment-related MDS and hemophagocytosis. Leukemia mutation panel was not performed. She ultimately passed due to disseminated candidemia in the setting of persistent pancytopenia.** | | | |

***FIGURES***

**Supplementary Figure S1. Box plots showing values of A) C-reactive protein (CRP), B) Ferritin and C) Lactate dehydrogenase (LDH) at baseline among patients with CH and those without CH.**

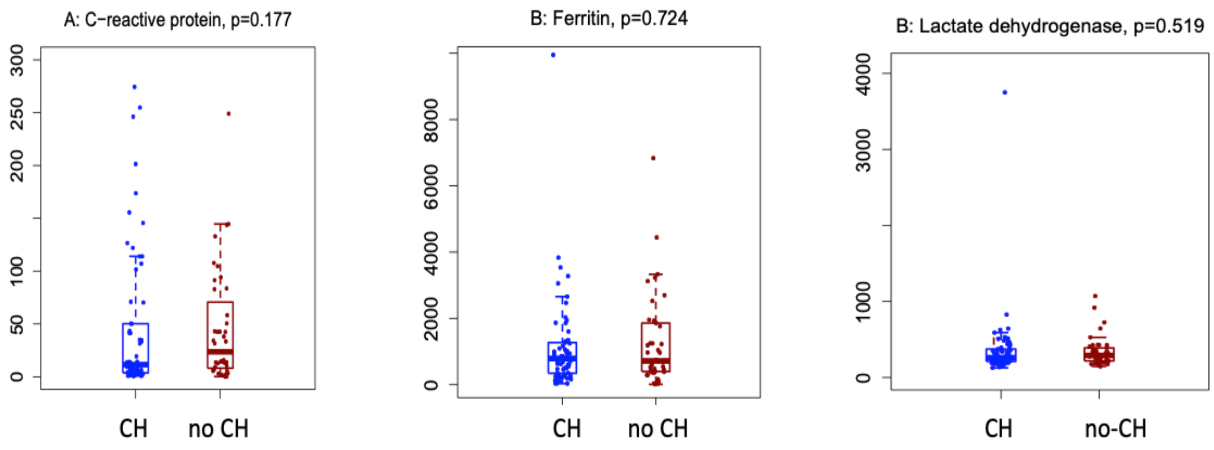


**Supplementary Figure S2. Incidence of different gene mutations implicated in clonal hematopoeisis in our cohort.**

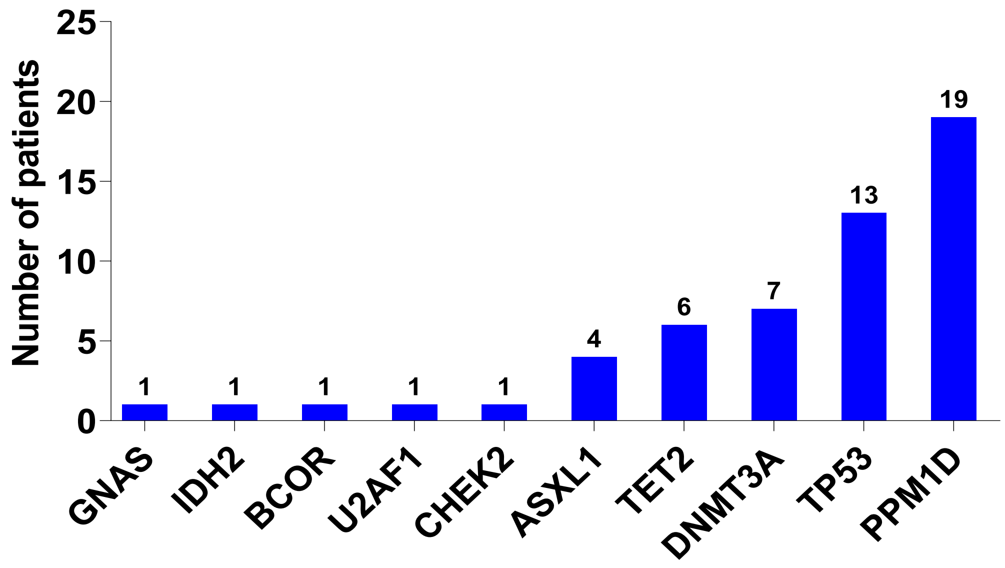

**Supplementary Figure S3. Bar graph plot showing median variant allele frequency of different CH mutations in our cohort.**

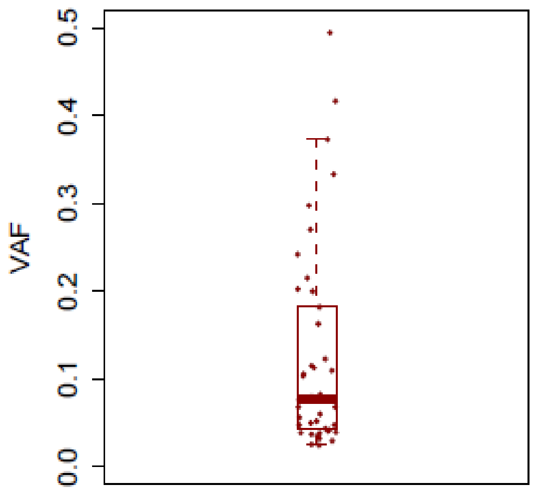


**Supplementary Figure S4. Kaplan-Meier Curve of the progression free survival (PFS) and overall survival (OS) of the whole cohort. The median follow-up time was 14.9 (range: 1.2-30.5) months. The median OS time was 15.7 (95%CI: 11.3-not reached) months, and the median PFS time was 4.8 (95%CI: 3.5-7.4) months.**

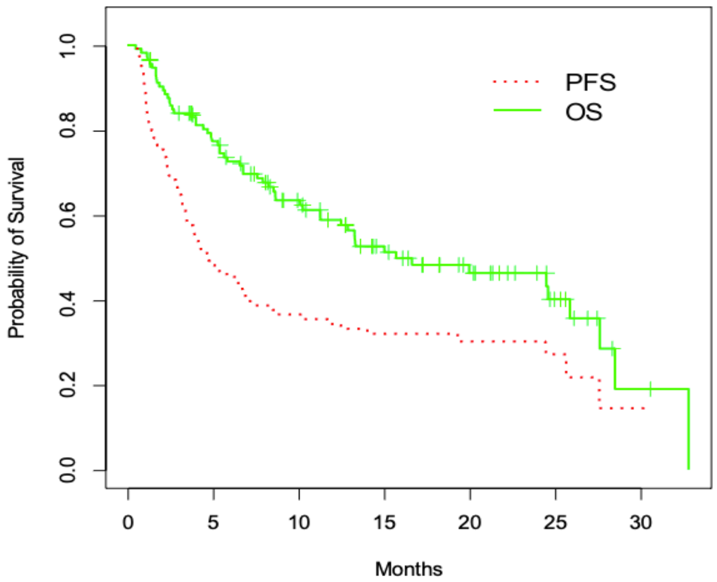

**Supplementary Figure S5. Kaplan-Meier Curve of overall survival, stratified by CH status. The p-value is 0.293 based on the log rank status.**

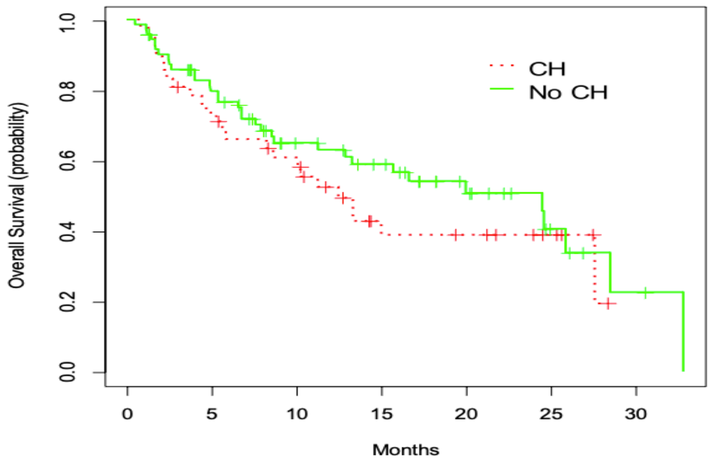


**Figure S6. Bar graph showing the percentages of patients requiring tocilizumab and steroids in the CH versus no-CH cohorts. The percentage of patients requiring tocilizumab and corticosteroids for management of CRS and ICANS was comparatively higher in patients with CH at 64.3% (27/42) and 52.4% (22/42), respectively, compared to 55.6% (40/72, p=0.43) and 43.1% (31/72, p=0.43) of patients with no-CH, respectively.**

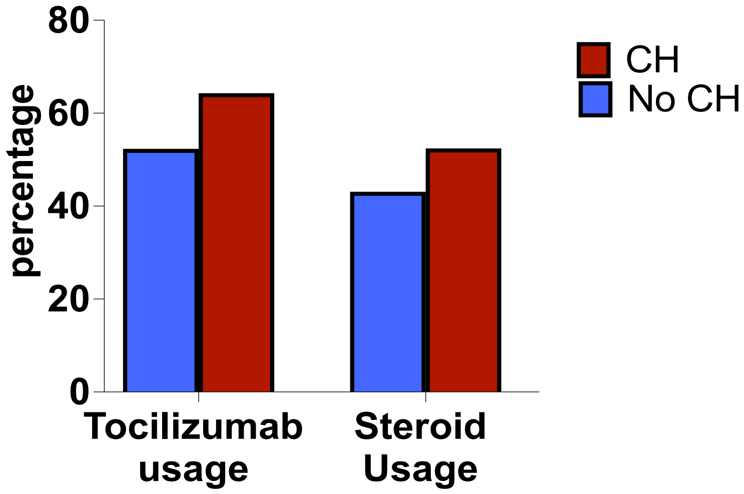
